# Respiratory and diarrhoeal pathogens in Malawian children hospitalized with diarrhoea and association with short-term growth

**DOI:** 10.1101/2021.10.13.464207

**Authors:** Mphatso Chisala, Wongani Nyangulu, James Nyirenda, Pui-Ying Iroh Tam

## Abstract

**Background:** Pneumonia and diarrhoea are the leading causes of childhood mortality and morbidity worldwide. Recurrence of these common infections are one of the immediate causes of malnutrition, which puts children at risk of further infection. While studies have focused on how gut microbiota is broadly protective against respiratory infection, there has been less attention paid to the reverse relationship, of respiratory microbiota and pathogens influencing the gut, and subsequent association with growth.

**Methods:** In this sub-study of a prospective cohort study, 27 children (2-24 months) who tested positive for *Cryptosporidium* were followed up over 8 weeks. Respiratory and stool pathogens were detected using quantitative molecular methods. Nutritional outcomes were assessed as length-for-age (LAZ), weight-for-length (WLZ) and weight-for-age (WAZ) z-scores. Changes over the study period were compared using repeated analysis of variance and mixed effects model analysis.

**Results:** In this period,104 sputum and stool samples were collected. All stool samples had at least one pathogen detected, with an average of 5.1 (SD 2.1) stool pathogens, compared to 84% of the sputum samples with an average 3.5 (SD1.8). Diarrhoeagenic *E. coli* were the most common stool pathogens (92%), followed by *Cryptosporidium* (52%) and *Campylobacter* pan (39%). In sputum, *S.pneumoniae* was most prevalent (84%), seconded by Rhinovirus (56%) and *M. catarrhalis* (50%). There was a significant change in WAZ over the follow-up period. Children who had ≥3 GI pathogens had significantly a lower LAZ mean score at enrollment (−1.8 (SD 1.4)) and across the follow-up period. No relationship between respiratory pathogens and short-term growth was observed. Out of 49 sputum samples that had ≥3 pathogens, 42 (85%) simultaneously had ≥3 GI pathogens.

**Conclusion:** Among young children hospitalized with diarrhoea, multiple gut and respiratory pathogens were prevalent over an 8-week follow-up period. The presence of more GI, but not respiratory, pathogens was significantly associated with reduced short-term growth.

**Author summary:** The gut-lung axis interact in both health and illness, and we aimed to see whether presence of pathogens in the GI and/or respiratory tract is associated with short-term growth. In 104 sputum and stool samples, we observed lower mean short-term growth in participants with higher number of GI, but not respiratory, pathogens.

## Introduction

Lower respiratory infections (LRI) and diarrhoeal diseases are the top leading preventable causes of mortality and morbidity globally in children under 5 years of age and are the annual cause of 12% (700,000) and 9% (446,000) deaths, respectively [1–3]. Children with frequent and recurrent infections are at risk of malnutrition which also predisposes them to further infection [4]. Malnutrition is one of the most important risk factors for both diarrhoeal disease and LRI [3,5], and is associated with about half of all under-5 deaths [6]. Reducing the burden of malnutrition therefore could concomitantly decrease respiratory and diarrhoeal disease amongst high-risk children. This was estimated in the Global Burden of Disease study to be a reduction of 9% of LRI and 12% of diarrhoeal diseases over the past three decades [1,3].

Recently, large scale studies including the “Etiology, Risk Factors and Interactions of Enteric Infections and Malnutrition and Consequences for Child Health and Development” (MAL-ED) study and the “Global Enteric Multi-site” (GEMS) study explored the association between gastrointestinal (GI) pathogens, malnutrition and gut function with other long-term effects to understand the pathophysiology of malnutrition [7–9]. These studies noted that subclinical infections and quantity of pathogens are negatively associated with linear growth, and that this persists till 2 years of age [8,10]. Cumulative insults of infections like diarrhoeal and LRI on children likely causes failure of catch-up growth resulting in growth faltering and decreased cognitive development [11–14]. However, how these pathogens are associated with other common infections seen in these children (including respiratory infections) and how they are associated with short-term growth has not been explored.

Both the gut and lungs have the same embryonic origin, and thus, while the mechanisms are not understood, studies have shown that the two sites interact in health and disease. Specifically, animal studies have demonstrated how host-associated gut and respiratory microbiota appears to influence local and systemic immunity [15–18]. This has been termed the ‘gut-lung axis’ [16]. However, while these studies have focused on how gut microbiota is broadly protective against respiratory infection [19], there has been less attention paid to the reverse relationship, of respiratory microbiota and pathogens influencing the gut, and subsequent association with growth. Studies have indicated a link between respiratory infections and growth [20,21], although in general the pathways between growth, nutrition and infection are likely bidirectional.

This objective of this study was to describe detection of diarrhoeal and respiratory infection in children who were hospitalized with diarrhoeal disease at a tertiary hospital in Malawi, and to examine the association with short-term growth over an 8 week follow up period.

## Methods

### Study design, setting, and participants

This is a secondary data analysis from a prospective longitudinal observational study evaluating respiratory cryptosporidiosis in pediatric diarrhoeal disease. Children presenting with primary gastrointestinal (GI) symptoms to Queen Elizabeth Central Hospital in Blantyre, Malawi, were enrolled in the study from March 2019 to April 2020. Children were eligible to participate if they were 2-24 months of age and had at least three or more loose stools within the past 48 hours and lived 15km outside of the study [22]. Children were excluded if they had visible blood in loose stools, or dysentery. Parents provided written informed consent. The study was approved by the University of Malawi College of Medicine Research Ethics Committee (P.07/18/2438) and the Liverpool School of Tropical Medicine Research Ethics Committee (18-066).

### Clinical procedures

Participants positive for *Cryptosporidium* in either respiratory or GI tract specimens at enrollment were followed up every 2-week until 8 weeks post-enrollment. At each visit, history and physical exam were conducted, and induced sputum and stool were collected. The induced sputum procedure has been previously described in detail elsewhere [22,23]. In brief, sputum samples were obtained via oropharyngeal suctioning after nebulized 3% sodium chloride treatment and processed for microscopy and multiplex PCR testing.

Diarrhoea was defined as ≥3 loose stools within the past 48 hours. A diarrhoeal episode was termed symptomatic if a stool sample collected was PCR positive for any pathogen and the participant reported any GI symptoms (to include abdominal pain/tenderness, dehydration, vomiting, and/or poor feeding) and asymptomatic if the stool was PCR-positive for any pathogen but the participant did not report any GI symptoms. If the respiratory specimen was PCR-positive for any pathogen and the participant reported any respiratory symptoms (to include cough, runny nose, difficulty in breathing, wheezing, chest indrawing/retractions, and/or crackles) then the episode was termed symptomatic, and asymptomatic if the specimen was PCR-positive in the absence of any reported respiratory symptoms.

Nutrition indices were defined according to WHO growth standards. Weight-for-length z score (WLZ) for wasting, weight-for-age z scores (WAZ) for underweight and length-for-age z scores (LAZ) for stunting with z-score <-2 defined as either being wasted, underweight or stunted, respectively.

### Laboratory procedures

We conducted DNA extraction from specimens using a QIAamp Fast DNA Mini Kit (Qiagen, Hilden, Germany) with a procedure modified from that of the manufacturer as previously described [24]. Appropriate negative and positive controls were included in every extraction and PCR run. Plate-based qPCR was performed as previously described [25]. These qPCRs were carried out using the ViiA7 Real-Time PCR instrument (Thermo Fisher, Waltham, MA, USA) or the QuantStudio 7 Flex Real-Time PCR instrument (Thermo Fisher).

In sputum, multiplex testing detected bacteria (*Streptococcus pneumoniae, Staphylococcus aureus, Moraxella catarrhalis, Bordetella pertussis, Haemophilus influenzae* and *Haemphilus influenzae* type b, *Chlamydia pneumoniae, Mycoplasma pneumoniae, Klebsiella pneumoniae, Legionella pneumophila*, and *Salmonella* species); viruses (influenza A/B/C, RSV A/B, parainfluenza virus types 1-4, coronaviruses NL63, 229E, OC43, and HKU1, human metapneumovirus A/B, rhinovirus, adenovirus, enterovirus, parechovirus, bocavirus, cytomegalovirus); and parasites (*Pneumocystis jiroveciĩ*). A cycle threshold (Ct) of <38 was considered as a positive for any of these pathogens. *Cryptosporidium* detection of respiratory specimens were measured using quantitative polymerase chain reaction (qPCR), with a positive result being a Ct of <35.

In stool, GI pathogens were detected using qPCR in a TaqMan Array Card (Thermo Fisher, Waltham, MA) using a custom design developed at the Houpt Laboratory (Charlottesville, VA [25]). Multiplex testing detected rotavirus, norovirus GII, adenovirus, astrovirus, sapovirus, enterotoxigenic *Escherichia coli* (ETEC), enteropathogenic *E. coli* (EPEC), enteroaggregative *E. coli* (EAEC), Shiga-toxigenic *E. coli* (STEC), Shigella/enteroinvasive *E. coli* (EIEC), *Salmonella, Campylobacter jejuni/coli, Vibrio cholerae, Clostridium difficile, Cryptosporidium, Giardia lamblia, Entamoeba histolytica, Ascaris lumbricoides*, and *Trichuris trichiura*). We considered a pathogen as present if Ct was <35 in all pathogens. For pathogens that were positive and are associated with diarrhoea in children under 5 years old [8,13], we calculated the prevalence of diarrhoeagenic Ct cut-offs (diarrhoea-associated Ct quantity) based on GEMS study to estimated prevalence of diarrhoeal samples in this population [25]. An increased pathogen load was defined as at least 3 pathogens present in each sample per participants per study visit to compare how these relate with other demographics [26].

### Statistical analysis

At enrolment, categorical variables were compared using Pearson’s X^2^ test or Fisher’s exact test. Continuous variables were compared using Student’s t tests or nonparametric Mann-Whitney U tests where data were nonnormally distributed. We presented diarrhoeagenic quantitative cut-off based on the GEMS study to estimate the burden of diarrhoea attributed to the common causes of diarrhoeal pathogens in this population. These cut-off values are useful in studies that have no controls or that do not have diarrhoea data like our study. Comparison for different characteristics across the 8-week study period was done using one-way repeated measures analysis of variance and mixed effect model analysis. Statistical significance was set at 0.05, characteristics that showed a significant change over the follow-up period were included in a mixed effect model analysis as confounders. Statistical analysis was performed using Stata software, version 16.

## Results

### Participants

From March 2019 to April 2020, 755 children were screened and 162 were recruited into the study. Of the 162 enrolled, 37 (23%) were positive for cryptosporidium, 36 were entered into follow-up, and 27 children (75%) completed the 8-week follow-up, which was discontinued early due to COVID-19. Of these, the median age was 5.5 (IQR 2,14) months and 18 (64%) were male. Only 1 (3%) was HIV-infected but HIV status was unknown in over half of the children (20/27). The enrolled study population is described elsewhere [23].

We tested 104 stool and sputum samples from the 27 participants that had completed follow-up. There was at least one pathogen detected in all the 104 stool samples while 87/104 (84%) of the sputum samples had at least one pathogen detected. Amongst the 104 stool samples collected, diarrhoeagenic *E. coli* were the most abundant pathogens (92/104 (89%)) of which EAEC was the most common subtype (88/92 (95%)) followed by typical EPEC (42/92 (45%)). *Cryptosporidium* species (55/104 (52%)) were the second most common stool pathogen followed by *Campylobacter* pan (41/104 (39%))*, Adenovirus* pan (34/104 (33%)) and Norovirus (27/104 (26%)). Over half (58/104 (56%)) of stool samples with a pathogen associated with diarrhoea were below the diarrhoeagenic cut-offs as presented in the GEMS study (Figure 1). Out of the 104 induced sputum samples, *S. pneumoniae* was the most abundant pathogen 87/104 (84%) followed by human Rhinovirus (58/104 (56%)), *M. catarrhalis* (52/104 (50%)), Adenovirus (29/104 (28%)) and *H. influenzae* (21/104 (20%)). The majority of stool samples (80/104 (77%)) and almost half of sputum samples (49/104 (47%)) had at least three pathogens (data not shown). Of the 49 sputum samples that had a high pathogen load, 42/49 (85%) simultaneously had ≥3 stools pathogens, although not statistically significant.

**Figure 1.**
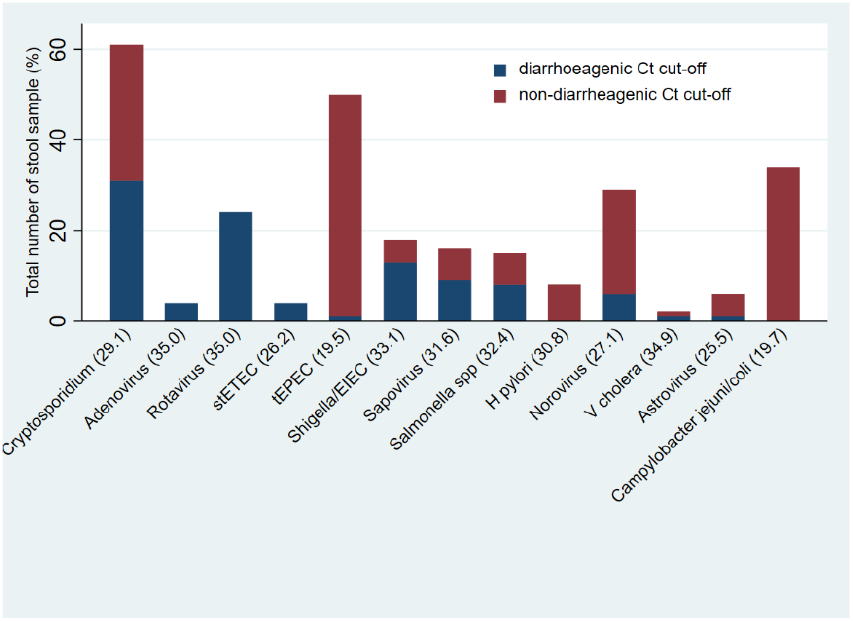
Positive stool samples with diarrhoeagenic cut-offs-as described in the GEMS study stETEC - heat-stable enterotoxin-producing *Escherichia coli*, tEPEC-typical enteropathogenic *Escherichia coli*, EIEC – enteroinvasive *Escherichia coli, H.pylori - Helicobacter pylori, V.Cholera-Vibrio Cholera*

GI pathogens were detected in all stool samples from 2 weeks to the end of 8 weeks. In contrast, respiratory pathogens were detected in all sputum samples at 2 weeks and in 20/25 (80%) samples the end of the 8 weeks. On average, there were 5.1 (SD 2.1) stool pathogens detected per participant per study visit over the follow-up period while an average of 3.5 (SD 1.8) sputum pathogens were detected per participant per study visit over the follow-up period (Table 1). There were an average of 3.1 (SD 1.6) bacteria compared to 1.0 (SD 0.8) parasites and 1.0 (SD 0.9) viruses in stool samples collected and an average of 2.0 (SD 1.1) bacteria, 1.4 (SD 1.1) viruses and 0.3 (SD 0.5) parasites in sputum samples (Figure 2).

**Table 1.**
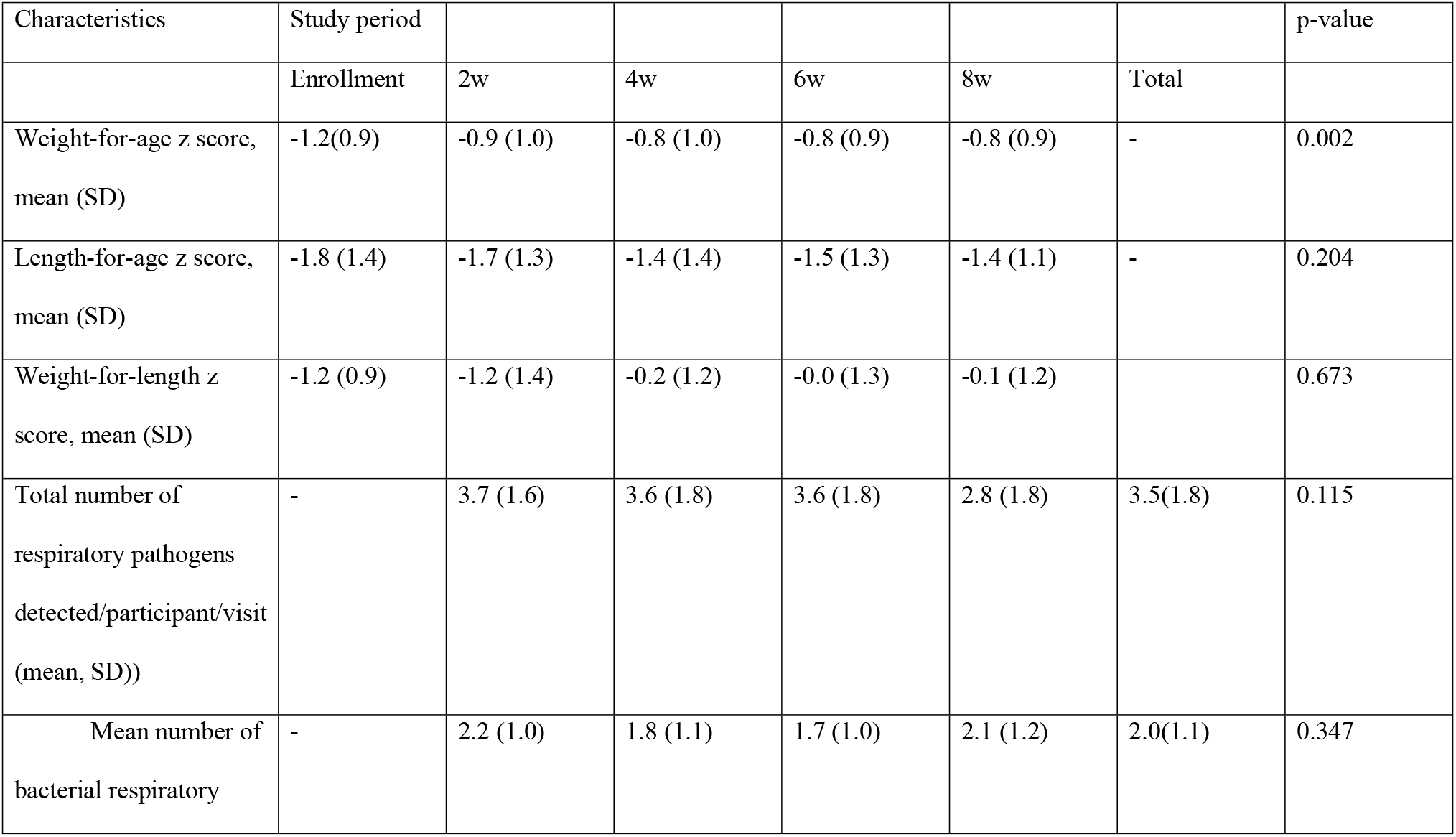
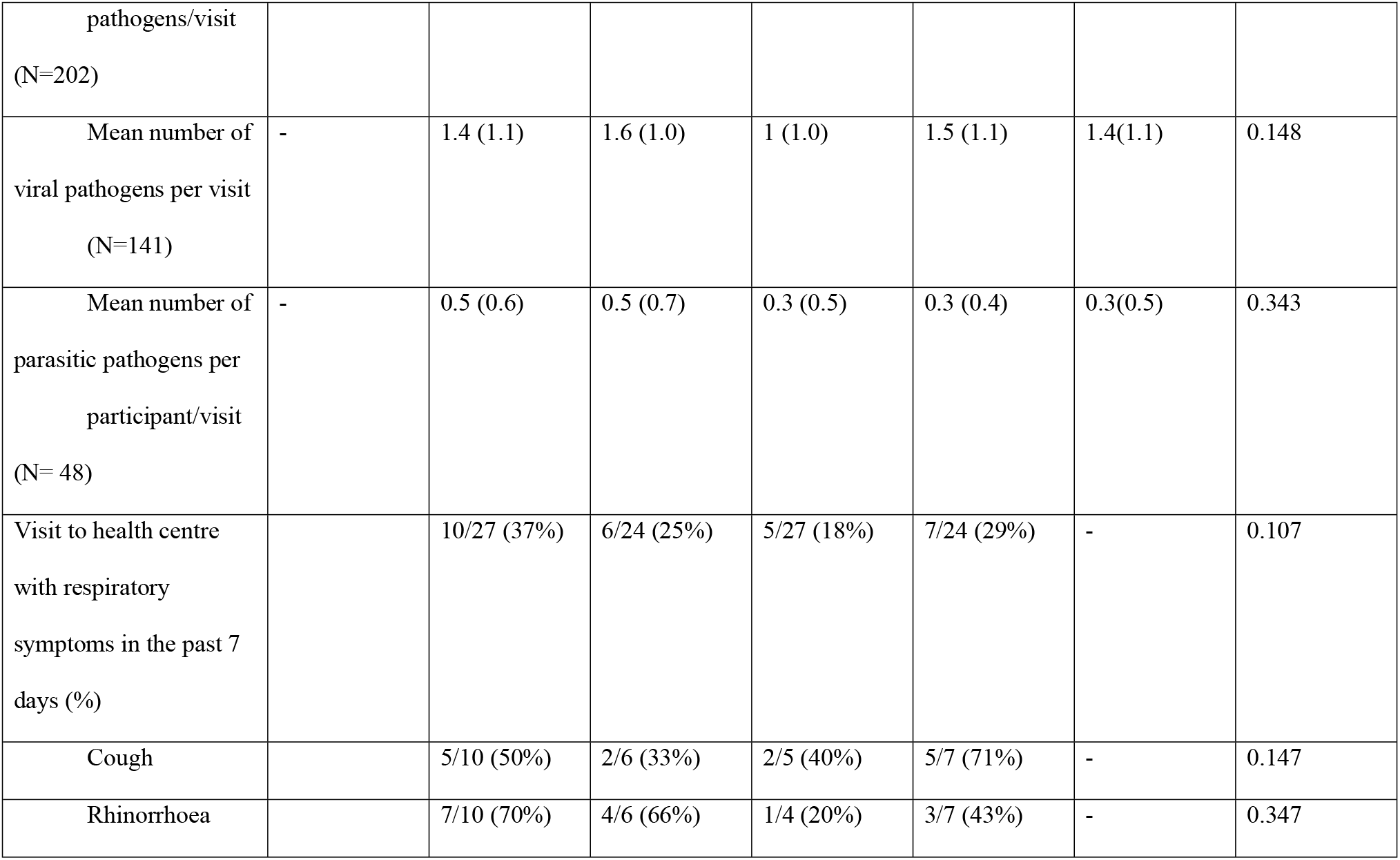
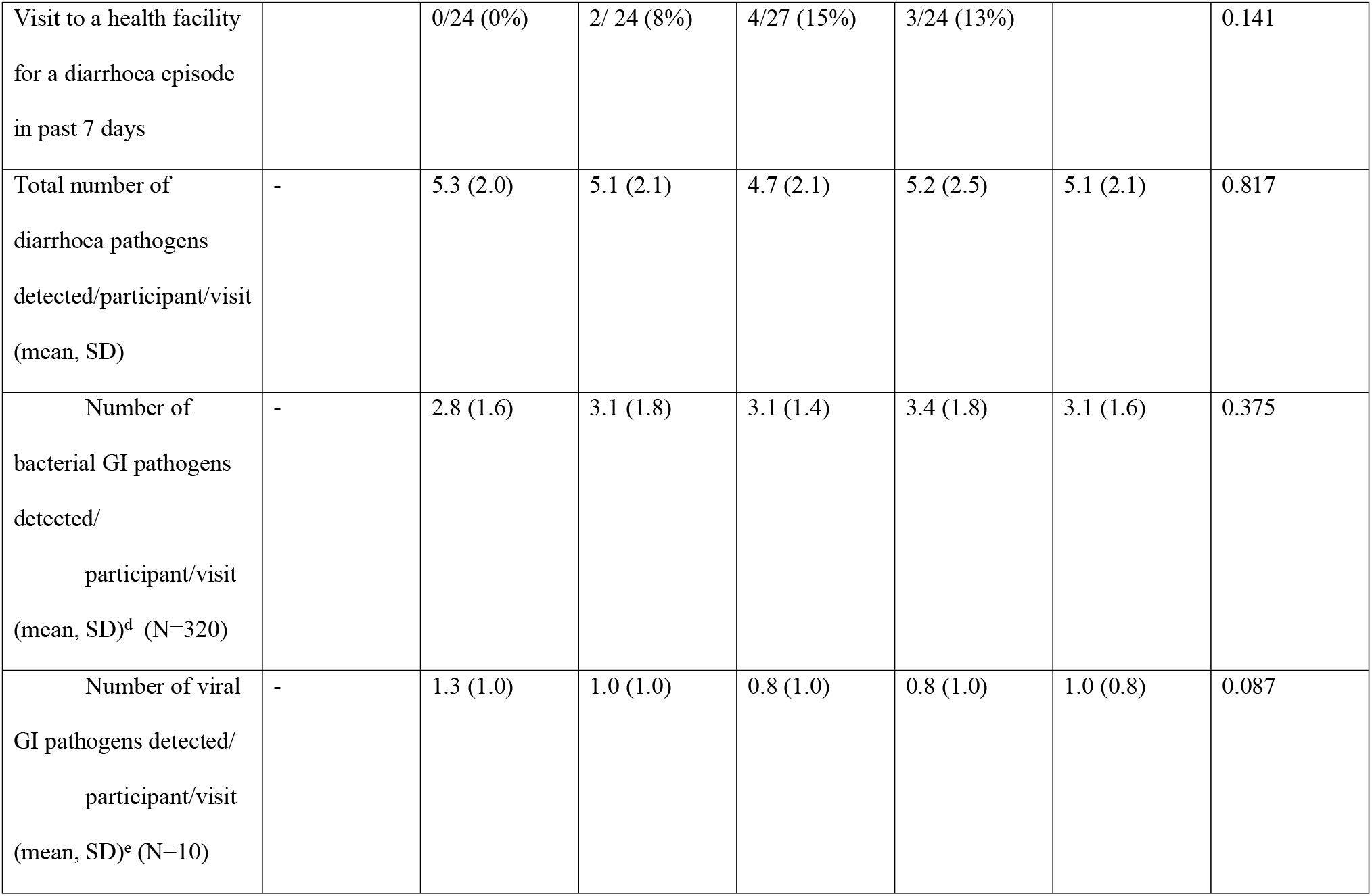
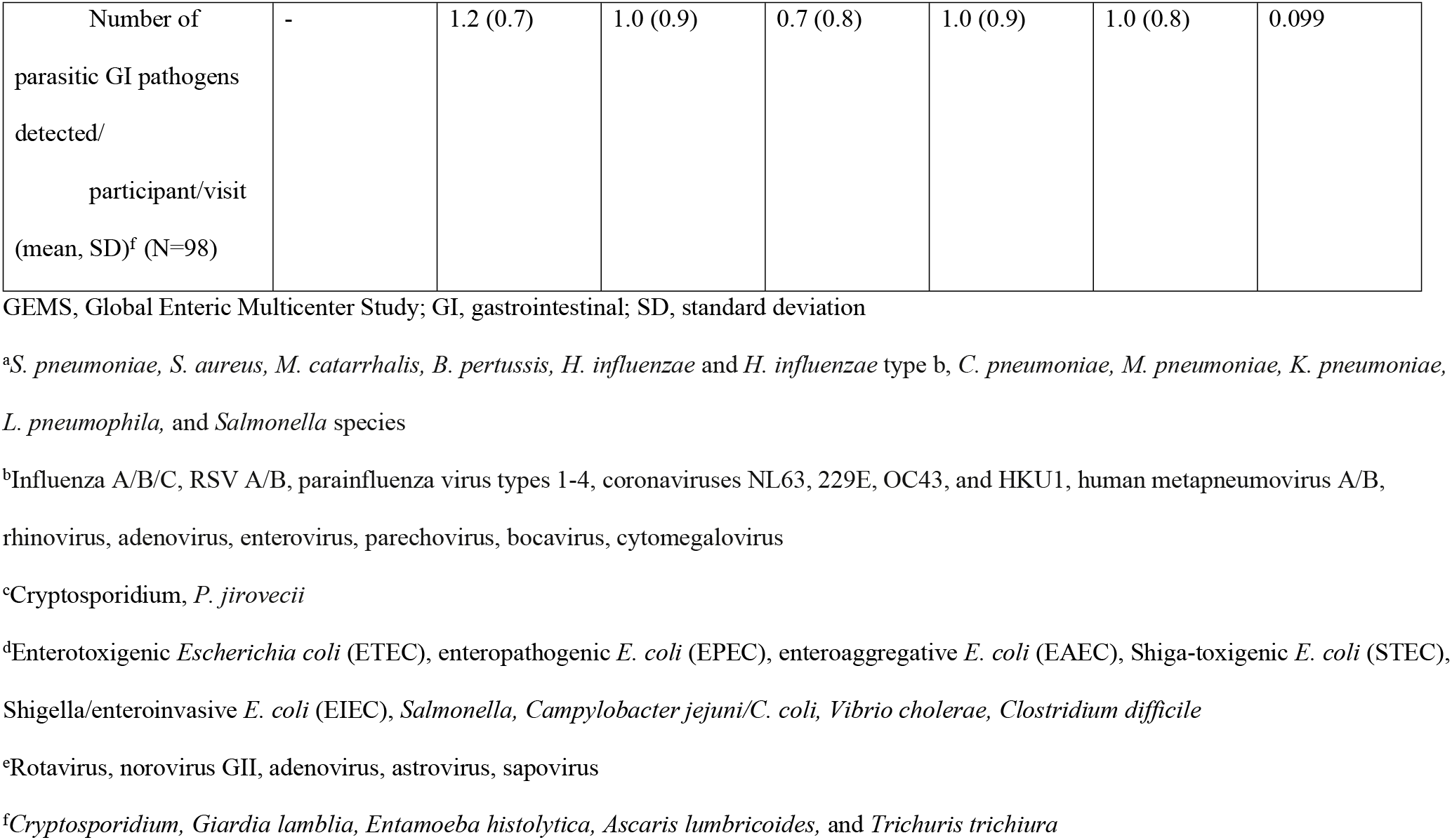
Characteristics of and change in study population over 8 weeks

**Figure 2.**
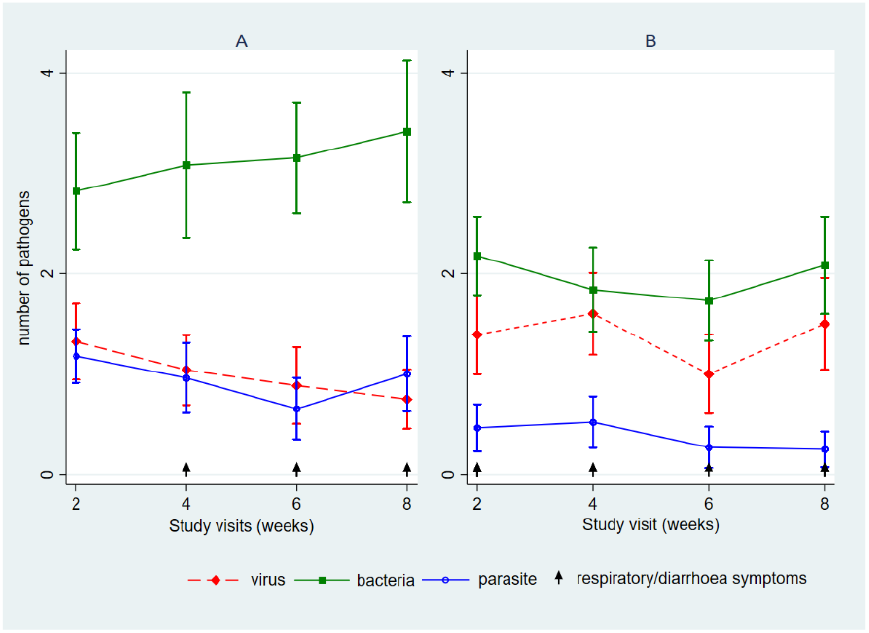
Average number of pathogens per study visit detected (mean, SD) over 8-week period in children with diarrhoeal disease in A) stool; B) sputum

Figure 3 shows the changes in WAZ, WLZ and LAZ across the study period. Participants had low LAZ and WAZ at enrolment, and the average change in WAZ, LAZ, and WLZ scores over the 8-week period were 0.5 (0.6), 0.4 (1.4) and 0.4 (1.4), respectively. There was a significant change in WAZ across the follow-up period (p-value-0.002), however no significant changes were seen in LAZ and WLZ. Children with ≥3 GI pathogens in a sample had lower LAZ compared to those with <3 pathogens at 2 weeks (−2.0±1.1 vs 0.1±0.5), 4 weeks (−1.6±1.4 vs −0.5±0.8), 6 weeks (−1.8±1.4 vs −1.0±1.2) and 8 weeks (−1.6±1.2 vs −0.6±0.9), and this was statistically significant(Figure 3B). This was not noted for WLZ and WAZ scores. No obvious changes in WLZ, WAZ and LAZ were noted with respiratory pathogen detection over the study period (Figure 3C). There was also no difference in any of the nutritional indices amongst children with ≥3 of both respiratory and gastrointestinal pathogen compared to those with <3 (Figure 3D).

**Figure 3.**
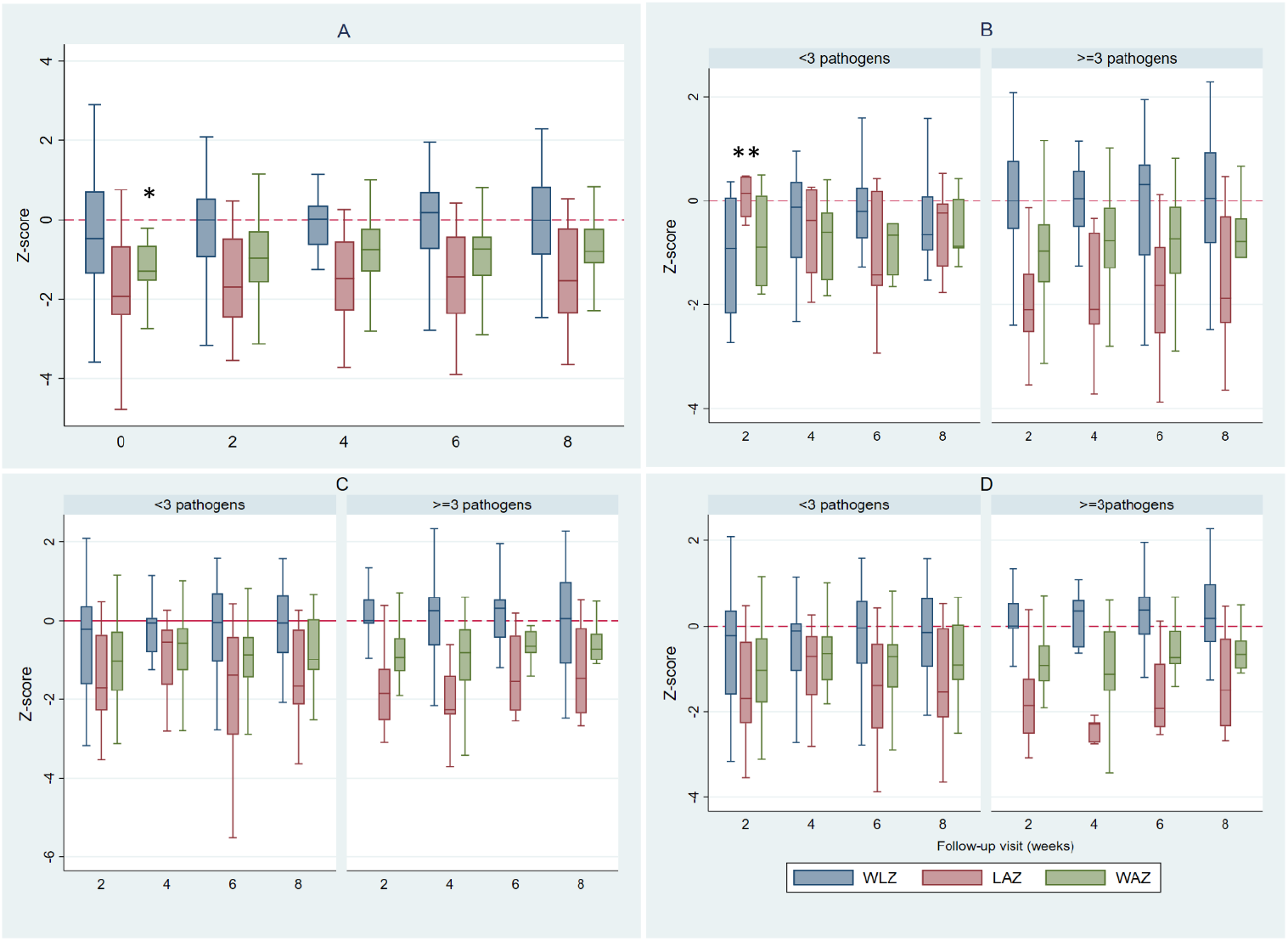
Change in mean WHZ, HAZ, LAZ scores over the 8-week study period: A) overall; B) in relation to average GI pathogens of i) <3 and ii) ≥3; C) in relation to average respiratory pathogens of i) <3 and ii) ≥3. D) in relation to respiratory and GI pathogens *p-value 0.006 **p-value 0.001

## Discussion

This is the first description, to our knowledge, of both respiratory and GI pathogens in young children and association with short-term growth in the 8 weeks after hospitalization with diarrhoea. We found that our population had low anthropometric indices, and that these indices showed minimal change over the 8 weeks after hospitalization. A high average number of GI pathogens was detected throughout the 8 weeks, and this was/not associated with gastrointestinal symptoms. A high average of respiratory pathogens was also detected throughout the 8 weeks, predominantly without associated respiratory symptoms. Significant changes were only noted in WAZ and not the other anthropometric measures over the 8-week follow-up period. Additionally, participants with ≥3 GI pathogens had a lower mean LAZ score at all follow-up visits.

Previous studies that have evaluated GI pathogens in young children after hospitalization and association with short-term growth have typically focused on a single pathogen. [12,27–30]. Short term growth after an infection in children under 2 years old is important because it allows for catch-up growth in this critical period for children to attain their optimal weight and height velocity, [11,31]. Similarly, in this same critical period, the gut microbiome is being established which is also associated with immune programming that affects long term health status within the gut and other distant organs like the lung [3].

Bacterial components and functions in the lung and gut can influence immune response and function in distal organs to influence health and disease, although mechanism are unclear [16]. This was also well demonstrated in a pre-clinical study by Wedgewood et al where pups exposed to malnutrition in the early days of life had an altered gut microbiota and developed hyperoxia and pulmonary hypertension which were corrected after ingestion of a probiotic [18]. The vast acute or ongoing immunological changes that happen in the gut as a result of insults like pathogens result into local, systemic, and distal organ changes, including inflammatory reactions to the lungs exposing them to acute or chronic inflammatory conditions [18,19]. These changes could equally disrupt commensal microbial balance in the lung and potentially putting them at risk of increased pathogenic microbial overgrowth and/or translocation. This could also explain why majority of participants with a high number of respiratory pathogens also had a high number of GI pathogens in our study.

Changes in the gut microbiota composition resulting from dietary components have been identified in chronic lung diseases and acute respiratory infections. For example, Santo et al noted that an increase in short chain fatty acids in the gut -from a high fiber diet-was strongly associated with reduced exacerbations of inflammatory lung diseases including asthma [33]. Collectively, studies reviewed by Criq et al in “dysbiosis, malnutrition and enhanced gut-lung axis contribute to age-related respiratory diseases,” highlighted that elderly people, who are often malnourished, present with a gut microbiota dysbiosis which likely plays a central role in the progression of ageing-related disorders including respiratory diseases, and that optimizing adequate nutrition amongst older people may be the most accessible lever of action to prevent the development of lung disease [34]. These studies demonstrate that there is an interaction between nutrition and the gut-lung axis.

The MAL-ED was a multisite cohort study where stool and length measurements were collected monthly from birth to age 2 years. Almost all (95%) stool specimens, from predominantly non-diarrhoeal episodes, detected at least one enteropathogen. Diarrhoea of any cause were associated with small reductions in LAZ after 3 months, and this difference was most pronounced when cause was attributed to bacteria, specifically EAEC, Campylobacter, and Shigella [35]. This aligns with our findings, where we identified an average of 5.1 (2.1) stool pathogens per stool sample, with diarrhoeagenic *E.coli* (particularly EAEC) being the most common enteropathogen. *Cryptosporidium* was prevalent throughout the follow-up period [23], and is similar to what was found in the GEMS and MAL-ED studies, which highlighted *Cryptosporidium* infection as a prevalent and persistent cause of diarrhoea amongst children <5 years, and associated with linear growth deficits [7,8].

In the Pneumonia Etiology Research for Child Health (PERCH) study, multiple respiratory pathogens were noted among children hospitalized with severe pneumonia, with an average of 3.8(SD 1.5) pathogens/participant [36]. However, these were participants with severe pneumonia and consecutive sampling was not done. Additionally, malnutrition was consistently more prevalent amongst these children with severe pneumonia (cases) compared to controls, ranging from 32% in Zambia [37] and 48% in Mali [38]. In our study, *S. pneumoniae* was detected in majority of cases, which is surprising given that we expect the prevalence to decrease in this setting where routine pneumococcal conjugate vaccination is available. We however did not analyze pneumococcal serotypes in our study which would also identify whether the prevalent *S.pneumoniae* strains are non-vaccine serotypes or residual vaccine serotypes as seen in similar settings [39]. *M. catarrhalis* was the most prevalent bacteria detected, with the most prevalent viruses detected being rhinovirus and adenovirus. It is also more likely that these respiratory microbes were colonizers and not pathogens of the respiratory tract, particularly as they were not associated with respiratory symptoms [40]. Malnutrition is a significant risk factor for respiratory infection, although the direct linkage of respiratory infection to growth is not fully explored. Hida et al noted that although there was no association between respiratory infections and growth, children who had respiratory infections and were supplemented with Vitamin A grew 0.17 cm/4 months (95% CI:0.002, 0.33) less than children with no respiratory infection who were treated with vitamin A, suggesting that respiratory infections may play an intermediate role in growth [20].

There was a high prevalence of stunting in this studying population. At enrolment, over a third of our study population was stunted and the mean LAZ was −1.8 (1.4) [23], similar to the national stunting prevalence amongst children <24 months [41]. We also noted that throughout the follow-up period, children who had a higher pathogen load were more stunted than those with a lower number of enteropathogen. These changes may be due to the high enteropathogen load, which are hypothesized to result in chronic gut inflammation and a leaky gut - an environmental enteropathy, a poorly understood, chronic condition seen in children from developing countries [42]. There was also a positive change in WAZ unlike WLZ and LAZ indices over the follow-up period. This would be because the children in thus population were more stunted at baseline and therefore could not see a change. Also important to note that although these children had diarrhoea at enrolment, the incidence of diarrhoea was much lower over the follow-up period. This positive change would therefore be as a result of catch-up growth (in weight) which is noted during a diarrhoea free period after an incident of diarrhoea [11,12,14]. We did not see an association between prevalence of respiratory pathogens with short-term growth, although we note that participants generally did not have respiratory symptoms. As demonstrated in the main CryptoResp study [23], presence of respiratory *Cryptosporidium* was significantly associated with gastrointestinal cryptosporidiosis. Similarly, majority of samples that had a high pathogen load in stools also had a high pathogen load in sputum. A healthy gut microbiota has been shown to broadly be protective of respiratory infections [16,19], although we cannot make any inferences or conclusions from this data.

Our study has limitations. The sample size was small, and not powered to detect changes in short-term growth. We did not have a comparison group of children with no *diarrhoea/Cryptosporidium* infection at baseline. We did not have stool or respiratory samples collected at the time of enrollment, and therefore could not assess change in prevalence of pathogens from time of hospitalization. However, these limitations are offset by the consecutive sampling of participants over an 8-week period from both respiratory and GI tracts, the collection of induced sputum from the respiratory tract, and the use of quantitative methods for analysis [24,40].

In summary, among young children hospitalized with diarrhoea, multiple gut and respiratory pathogens were prevalent in the participants over the following 8 weeks, and the presence of more GI pathogens, but not respiratory pathogens, was associated with reduced short-term growth. Further study of larger cohorts are warranted, to delineate how gut and respiratory pathogens interact and contribute to linear deficits, during a period where insults that occur can result in long term deranged growth, developmental and cognitive outcomes [32,43].

## Acknowledgements

We thank the patients and parents for participating in this study. We thank the research clinical and laboratory staff for conducting the study. We thank David Moore and Tanja Adams with training the CryptoResp clinical and laboratory teams. We thank Neema Toto for interim study support, Thokozani Ganiza for assistance with data management, and Wes Van Voorhis for reviewing a draft of the manuscript and providing critical feedback.

## Author contributions

Conceived and designed the study: MC, PI; Performed the study: MC, WN, HT, JN; Analyzed the data: MC, JN; Supervised the study: MC, WN, PI; Wrote first draft of the manuscript: MC, PI; Reviewed, provided critical feedback, and approved the final draft: all authors.

